# Single-cell Transcriptome-Proteome-PTMome(×8)-Metabolome (scTPPM) co-profiling decodes multimodal regulation during mouse oocyte meiosis

**DOI:** 10.64898/2026.11.09.765486

**Authors:** Yi-Rong Jiang, Lan-Rui Cao, Xiao-Feng Ni, Jie Wu, Yi Yang, Qin-Qin Xu, Yu-Meng Qiu, Ya-Xue Hou, Qiong Wu, Yu Wang, Hong-Shan Guo, Jian-Zhang Pan, Xu-Dong Fu, Qun Fang

**Author notes:** These authors contributed equally: Yi-Rong Jiang, Lan-Rui Cao, Xiao-Feng Ni, Jie Wu.

## Abstract

Single-cell multi-omics technology is crucial for comprehensive characterization of cellular molecular features. Despite recent advances in single-cell multi-omics approaches, co-profiling of nucleic acids, proteins, metabolites, and post-translational modifications (PTMs) within the same cell individuals remains major challenges. Here, we developed a single-cell transcriptome, proteome, PTMome, and metabolome (scTPPM) profiling approach, enabling simultaneous analysis of transcriptomes, proteomes, eight types of PTMs (e.g., phosphorylation, methylation, ubiquitination, O_GlcNAc, acetylation, crotonylation, lactylation, and succinylation), and metabolomes in individual cells. We demonstrated the scTPPM’s performance in tumor cell lines and oocyte samples, reaching total identification and quantification of 27,668 genes, 4,847 proteins, 1,508 PTM sites, and 680 metabolites in single mouse oocytes. We profiled previously-uncharacterized dynamic crosstalk among the entire metabolic pathways from metabolic enzyme expression and regulation to metabolite-mediated post-translational modifications during oocyte maturation. In summary, scTPPM provided an effective workflow for multi-omics co-profiling in single cells, enabling multi-layered insights into complex regulatory network.

## Main

Single-cell multi-omics technologies provide a systematic perspective for understanding cellular heterogeneity, offering remarkable advantages in revealing the mechanisms of complex biological processes^1,2^. Beyond transcriptional regulation, emerging evidence highlights the critical roles of protein post-translational modifications (PTMs) and metabolic reprogramming in shaping cellular phenotypes^3,4^, especially in the formation of cellular dynamic regulatory network during development and disease pathogenesis. To further capture these complex interactions, advanced technologies capable of simultaneously profiling transcriptomes, proteomes, PTMomes, and metabolomes at the single-cell level are urgently required.

The molecular components in single cells include nucleic acids, proteins, and metabolites. Simultaneous analysis of all these three major categories of components within individual cells presents fundamental technical challenges, primarily due to the diverse nature of these components, incompatibility among different analysis platforms, and sensitivity limitations. In particular, the initial quantities of the components within single cells are extremely low, and proteins and metabolites cannot be amplified like nucleic acids. Existing single-cell multi-omics approaches typically focus on nucleic acids, enabling simultaneous analysis of the genome, epigenome, transcriptome, and/or translatome^5–11^. In addition, several single-cell multi-omics approaches based on the microfluidics technique can achieve simultaneous analysis of nucleic acids and proteins^12–22^, while cannot accommodate metabolome analysis. Furthermore, the single-cell multi-omics approaches based on mass spectrometry can achieve simultaneous analysis of proteins, metabolites, and/or post-translational modifications (PTMs)^23–28^, but cannot accommodate nucleic acid sequencing. In summary, the current single-cell multi-omics approaches cannot achieve simultaneous analysis of nucleic acids, proteins, and metabolites, let alone combining the analysis of PTMs, which presents even greater challenges.

Recently, the multi-omics approaches for bulk samples have enabled the integrated analysis of the transcriptome, proteome, PTMome, and metabolome^29,30^, primarily through sample homogenization and splitting followed by separate omics profiling. However, performing such strategies at the single-cell level remains a significant challenge, due to the extremely low abundance of target compounds, low reaction efficiency, complicated operation procedures, and substantial sample losses by non-specific adsorption. These challenges significantly hinder the implementation of multi-omics technologies enabling simultaneous profiling of nucleic acids, proteins, PTMs, and metabolites at the single-cell level.

In this study, we developed scTPPM, a novel microfluidics-based approach that leverages a precise nanoliter-scale sample-splitting strategy and highly sensitive subcellular multi-omics analysis approaches to simultaneously profile the transcriptome, proteome, PTMome, and metabolome from the same single-cell individuals. Based on the high-quality subcellular proteomic data and a novel multi-PTM library, we achieved in-depth analysis of an unprecedented eight types of PTMs from single cells with over a thousand of PTM sites. The scTPPM approach was applied to uncover dynamic variations in transcriptional, translational, PTM, and metabolic regulatory networks during mouse oocyte meiosis at the single-cell resolution. The obtained comprehensive datasets provided previously-uncharacterized features involving metabolite-mediated post-translational modifications and multimodal dynamic heterogeneity during oocyte maturation. The scTPPM approach enables simultaneous analysis of transcriptomics, proteomics, PTMomics, and metabolomics in single cells, as well as the complex correlations among these four layers of regulations, offering a new tool for characterizing single-cell intermolecular regulatory systems.

## Results

### Development and validation of scTPPM approach

Cellular activities are regulated by complex and dynamic interactions among nucleic acids, proteins, and metabolites. Therefore, simultaneous analysis of these biomolecules is crucial for accurately capturing the physiological state of individual cells. However, the challenge in developing this simultaneous analysis techniques arises from the fundamental incompatibility of the standard approaches for analyzing each family of biomolecules. Specifically, nucleic acids are primarily analyzed using next-generation sequencing techniques, proteins are mainly analyzed using antibody labeling or mass spectrometry-based approaches, and metabolites are predominantly analyzed using mass spectrometry methods.

To overcome the methodological incompatibilities mentioned above, we adopted a nanoliter-scale microfluidic sample-splitting strategy, which involves uniform single-cell lysis and lysate splitting for separate analysis of nucleic acids, proteins, PTMs, and metabolites. The conventional microliter-scale sample splitting methods are often ineffective for multi-omics analysis of single-cell samples, primarily due to the reduced reaction efficiency and increased non-specific adsorption caused by excessive dilution. To address this issue, the sample reaction volumes should be drastically reduced to the nanoliter scale, which significantly increases the difficulty of sample splitting and poses greater challenges. In this work, we utilized the microfluidic sequential operation droplet array (SODA) technique^31,32^ previously developed by the authors’ group capable of performing precise manipulation of liquids ranging from nanoliters to picoliters, to achieve precise and robust sample splitting at the hundreds-of-nanoliter scale for single-cell multi-omics analysis.

However, the use of the single-cell sample splitting strategy further reduced the sample amounts available for individual omics analysis from intact single cells to subcellular levels, posing greater challenges to the sensitivity of each omics analysis. Consequently, we optimized and improved the respective multi-omics analysis methods to accommodate the subcellular-level requirements. Specifically, in transcriptome sequencing, we reduced the reaction volume of the reverse transcription of transcripts from tens of microliters to hundreds of nanoliters, which significantly enhanced the depth of transcriptome sequencing. For proteome analysis, we used the nanoliter-scale systems for protein sample pretreatment to significantly increase the identification depth for subcellular-level samples. Moreover, the combination of nanoflow liquid chromatography and ion mobility enrichment significantly improved the analytical sensitivity in single-cell metabolomics.

In the terms of PTMs, conventional PTM analysis requires the enrichment of specific modifications from bulk samples prior to mass spectrometry-based identification and quantification. However, the enrichment of even one type of PTM from single-cell samples is challenging, let alone multiple types of PTMs. Recently, single-cell PTM analysis without enrichment has been reported with limited types of PTMs^33,34^, including phosphorylation, acetylation, methylation, or glycosylation. When attempting to further increase the number of PTM types analyzed simultaneously, the current analytical pipeline revealed its limitations in depth and efficiency. To address this, we developed a workflow to establish a multi-PTM library for library-based analysis of data-independent acquisition (DIA) data. This provides a novel strategy for exploring deeper information of PTMomes from single-cell data for further biological insights.

Based on the mentioned approach design, we established a workflow for single-cell Transcriptome-Proteome-PTMome-Metabolome (scTPPM) profiling (Fig. 1a). The performance of the scTPPM workflow was demonstrated in tumor cell lines and oocyte samples, reaching a simultaneous total identification of 9,592 genes, 2,519 proteins, 180 PTM sites, and 279 metabolites in single HeLa cells, as well as 16,221 genes, 3,885 proteins, 814 PTM sites, and 628 metabolites in single mouse oocytes (Fig. 1b). The results of the validation experiments indicated that the identification and quantitative results obtained using the partial samples from sample splitting showed no significant difference compared to the total single-cell samples (Fig. 1c, d). In addition, we compared our data with the published data^35,36^, which also demonstrated good consistency and correlation (Supplementary Fig. S1). Furthermore, a benchmark analysis was conducted to evaluate the quantitative accuracy of proteomic data obtained by scTPPM, and the standard error between the measurement values and the theoretical calculated values was less than 10% (Supplementary Fig. S2).

**Fig. 1:**
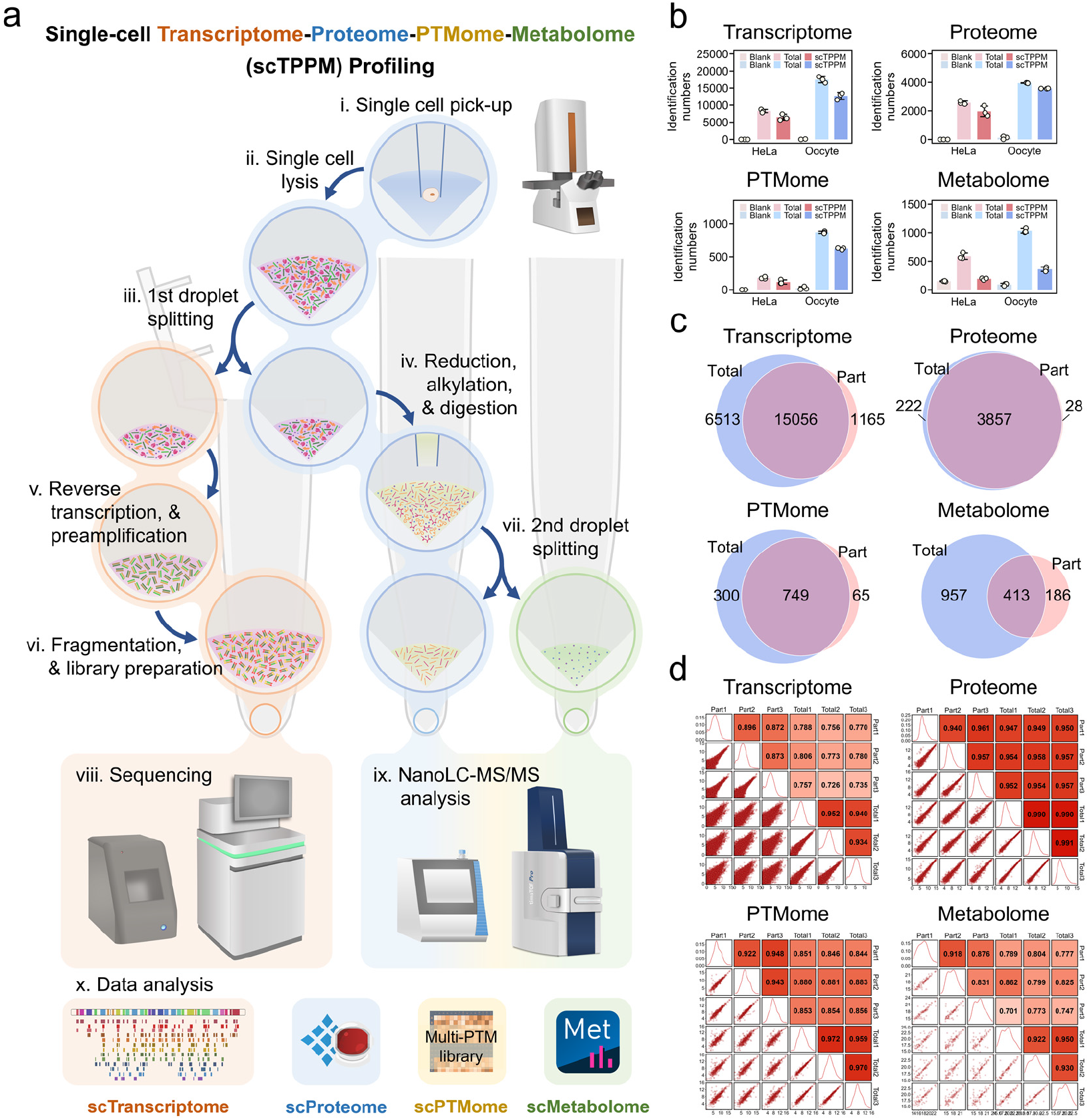
Development and validation of the scTPPM approach. a, The workflow of the scTPPM approach. b, The identification numbers of genes, protein groups, PTM sites, and metabolites in total (n = 3 for each omics) and partial (n = 3 for each sample type) samples of the single HeLa cells and single mouse oocytes. c, The Venn diagrams of the identified genes, protein groups, PTM sites, and metabolites between the total and partial samples of the single mouse oocytes. d, The correlation analysis of the quantified genes, protein groups, PTM sites and metabolites between the total and partial samples of the single mouse oocytes.

### Transcriptome, proteome, multi-PTMomes, and metabolome co-profiling during mouse oocyte maturation at the single-cell level

The scTPPM platform and workflow was applied to profiling the single-cell multi-omics of mouse oocytes during meiosis maturation (Fig. 2a), obtaining the expression information of transcriptome, proteome, multi-PTMomes, and metabolome in single oocyte samples at GV, MI, and MII stages (n = 8 for each stage). This dataset covers 27,668 transcribed genes (Fig. 2b), 4,847 protein groups (Fig. 2c), and 680 metabolites (Fig. 2d) in single oocyte samples. Using the unified symbol format, the genes identified in the transcriptome covered 98% of the genes identified in the proteome (Fig. 2e). In addition, we employed the self-built multi-PTM library to analyze eight types of PTMs (Fig. 2f, and Supplementary Fig. S3), identifying 465 phosphorylation sites, 413 methylation sites, 153 ubiquitination sites, 95 N-acetylglucosamylation sites, 94 acetylation sites, 91 crotonylation sites, 98 lactylation sites, and 99 succinylation sites (Fig. 2g). In the PCA dimension reduction and KNN cluster analysis, the oocyte samples of three different meiotic stages could be divided into three distinct clusters, especially using the proteome and PTMome data (Supplementary Fig. S4a). Fuzzy C-Means clustering of the transcriptomic data was able to categorize the genes into four clusters (C1-C4) with different expression patterns (Supplementary Fig. S4b). GO enrichment analysis further indicated that C1 and C4 were enriched in the RNA metabolism and phospholipid metabolic processes, C2 was related to ribosomal protein biosynthesis and chromosome segregation, and C3 was related to histone modification and cell cycle regulation (Supplementary Fig. S4c). Differential expression analysis of the proteomic data identified 287 upregulated proteins and 569 downregulated proteins from the GV to MII stage (Supplementary Fig. S4d). Differential expression analysis of the metabolomic data indicated that the oocytes at MI stage exhibit significant variation, and 20 differentially abundant metabolites were identified between the MI and GV samples (Supplementary Fig. S4e).

**Fig. 2:**
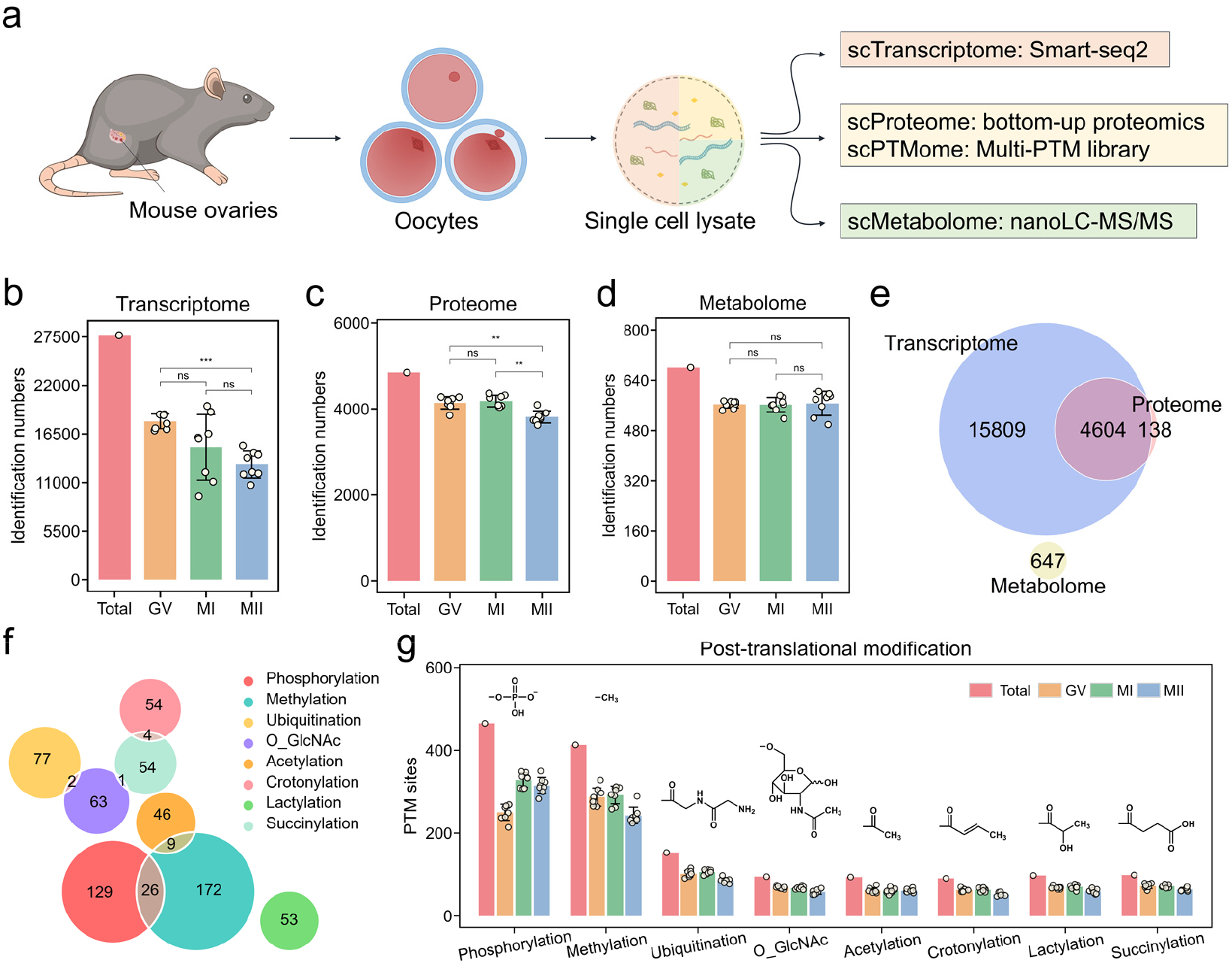
Transcriptome, proteome, multi-PTMomes, and metabolome co-profiling of single mouse oocytes at different meiotic stage. a, Schematic diagram of single-cell multi-omics analysis of single mouse oocytes. b, Identification numbers of transcribed genes in the GV (n = 8), MI (n = 8), and MII (n = 8) samples. c, Identification numbers of protein groups in GV (n = 8), MI (n = 8), and MII (n = 8) samples. d, Identification numbers of metabolites in the GV (n = 8), MI (n = 8), and MII (n = 8) samples. e, Venn diagram of the total transcribed genes, protein groups, and metabolites in single mouse oocytes (n = 24). f, Set relationships among eight types of PTMs using Euler diagram. g, Identification numbers of eight types of PTM sites in the GV (n = 8), MI (n = 8), and MII (n = 8) samples. *, p < 0.05; **, p < 0.01; ***, p < 0.001; ns, not significant; assessed by the student *t*-test.

### Multiple PTM sites with differential abundance identified in the subcortical maternal complex and cytoplasmic lattices

We further performed differential expression analysis of the PTM sites in the single mouse oocyte samples across the three meiosis stages. From the GV stage to the MII stage, a total of 527 differentially expressed post-translational modification sites were identified (Fig. 3a), covering 99 phosphorylated proteins and 89 methylated proteins (Fig. 3b). In the set relationship analysis, we observed an enrichment of differentially expressed PTM sites on key proteins of the subcortical maternal complex (SCMC)^37^ and cytoplasmic lattices (CPLs)^38^, such as Tle6, Nlrp4f, Khdc3, Nlrp5, and Padi6 (Fig. 3b), which played crucial roles in oocyte maturation and early embryonic development^39^. The prediction of the spatial distribution of PTM sites on the SCMC revealed that these PTM sites are located on the surface regions of the complex (Fig. 3c), indicating their functional relevance and accessibility. Notably, a significant upregulation of several PTM sites was identified at the MII stage, including the phosphorylation modifications of Nlrp5 and Tle6, as well as the phosphorylation, methylation, and glycosylation modifications of Khdc3 (Fig. 3d), which was consistent with the significant upregulation of Mad2l1 recruited by Khdc3 at the MII stage (Fig. 3e). Since Mad2l1 is an important spindle assembly checkpoint protein^40^, it is suggested that PTM sites on the SCMC are closely associated with the regulation of the oocyte meiosis. Furthermore, the protein expression levels of these complexes showed no significant upregulation, or were even significantly downregulated at the MII stage, such as Nlrp5, accompanied by an increase in ubiquitination levels (Fig. 3f). This result indicates the crucial and specific role of PTM regulation in the oocyte meiosis, independent of the transcriptional and translational processes.

**Fig. 3:**
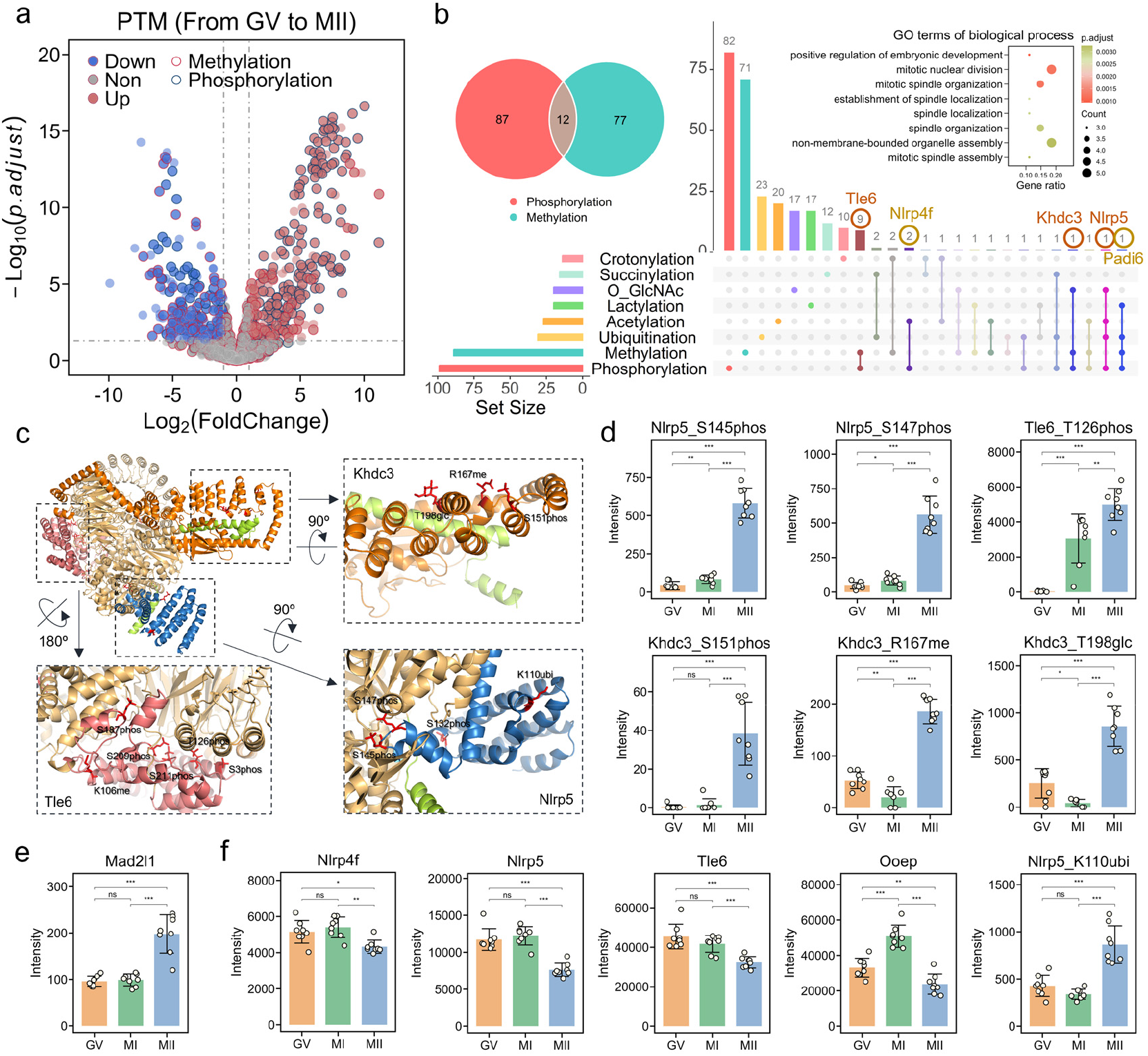
Deep multi-PTMomes profiling of single mouse oocytes during meiosis maturation. a, Differential expression analysis of PTM sites in mouse oocytes from the GV (n = 8) stage to the MII (n = 8) stage. b, Set relationships among proteins with differentially abundant PTM sites using Venn and Upset diagram, and GO enrichment analysis of proteins with more than two differentially abundant PTM sites. c, Spatial prediction of proteins and PTM sites in the SCMC using the published protein structure and AlphaFold3. d, Expression patterns of phosphorylation of Nlrp5, Tle6, and Khdc3, methylation and glycosylation of Khdc3 at the GV (n = 8), MI (n = 8), and MII (n = 8) stages. e, Expression pattern of Mad2l1 in mouse oocytes at the GV (n = 8), MI (n = 8), and MII (n = 8) stages. f, Expression patterns of Nlrp4f, Nlrp5, Tle6, Ooep, and ubiquitination of Nlrp5 in mouse oocytes at the GV (n = 8), MI (n = 8), and MII (n = 8) stages. *, p < 0.05; **, p < 0.01; ***, p < 0.001; ns, not significant; assessed by the student *t*-test.

### Histone deacetylase-mediated histone deacetylation represents a characteristic epigenetic feature during oocyte maturation

Histone modification regulation is one of the key mechanisms in the cellular epigenetic regulation. Using the scTPPM approach, we simultaneously identified six acetylation sites (H2K6ac, H3K18ac, H3K23ac, H3K27ac, H4K12ac, and H4K16ac), three methylation sites (H3K36me, H3K37me, and H3K79me), and one phosphorylation site (H3Y41phos) of histone in the single mouse oocytes (n = 24). During the oocyte maturation, although the expression levels of the histones remain relatively stable, the level of histone acetylation exhibited a significant downregulation from GV stage to MI stage for all six acetylation sites (Fig. 4a). This phenomenon was further supported by the immunofluorescence results (Fig. 4b, c), which is also consistent with the previous report^41^. Histone deacetylation can be defined as a significant epigenetic regulatory event during oocyte maturation, which regulates the spindle assembly and meiotic progression through its impact on chromatin accessibility^42,43^. Inadequate histone deacetylation can contribute to meiotic errors, aneuploidy, and abnormalities in early embryonic development^44,45^.

**Fig. 4:**
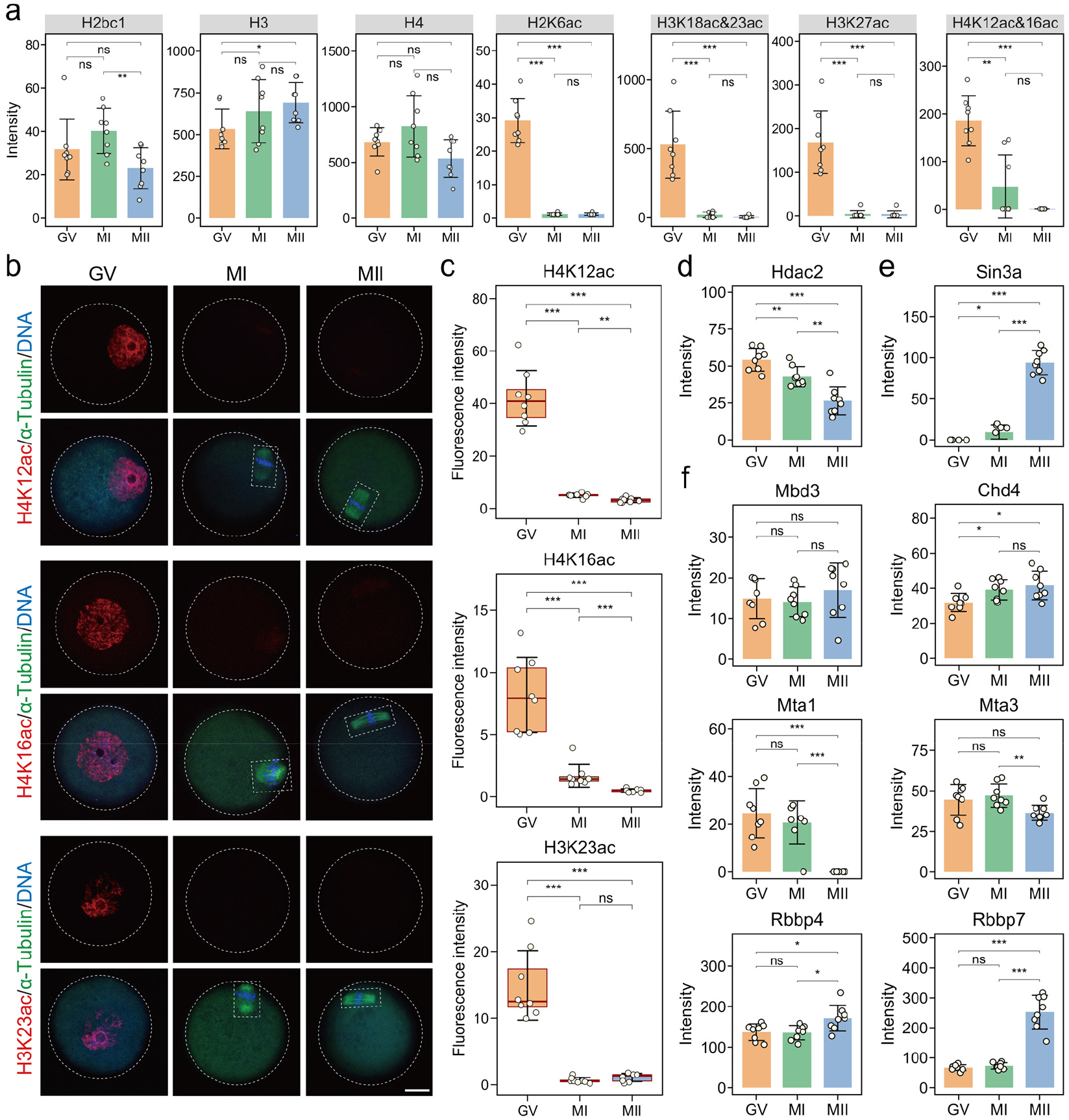
Histone acetylation and deacetylation of single mouse oocytes during meiosis maturation. a, Expression patterns of histone (H2bc1, H3, and H4) and histone acetylation (H2K6ac, H3K18ac, H3K23ac, H3K27ac, H4K12ac, and H4K16ac) in mouse oocytes at the GV (n = 8), MI (n = 8), and MII (n = 8) stages. b, Mouse oocytes at the GV, MI, and MII stages were immunostained on acetyl histone (H3K23ac, H4K12ac, and H4K16ac; Alexa Fluor 647, red) and α-tubulin (FITC, green), and fluorescently stained on DNA (DAPI, blue), scale bar: 25 μm. c, Fluorescent quantitative results of oocytes immunostained on acetyl histone (H3K23ac, H4K12ac, and H4K16ac; Alexa Fluor 647, red) at the GV (n = 8), MI (n = 8), and MII (n = 8) stages. d, Expression patterns of Hdac2 in mouse oocytes at the GV (n = 8), MI (n = 8), and MII (n = 8) stages. e, Expression patterns of Sin3a in mouse oocytes at the GV (n = 8), MI (n = 8), and MII (n = 8) stages. f, Expression patterns of Mbd3, Chd4, Mta1, Mta3, Rbbp4 and Rbbp7 in mouse oocytes at the GV (n = 8), MI (n = 8), and MII (n = 8) stages. *, p < 0.05; **, p < 0.01; ***, p < 0.001; ns, not significant; assessed by the student *t*-test.

Histone deacetylases (HDACs) are key erasers of histone acetylation, which can be divided into four classes^46^. Based on the multi-omics dataset, Hdac2 exhibits high expression abundance and an expression pattern consistent with the histone deacetylation process (Fig. 4d). Therefore, we propose that Hdac2 serves as the primary histone deacetylase during oocyte meiosis. The deacetylase function of Hdac2 is typically mediated within multi-subunit complexes^47^, primarily the Nucleosome Remodeling and Deacetylase (NuRD) complex^48^ and the Sin3 complex^49^. Notably, Sin3a, a core component of the Sin3 complex, exhibits extremely low abundance at the GV stage and the MI stage (Fig. 4e). Since the histone deacetylation occurs from the GV stage to the MI stage, we propose that the NuRD complex may play a crucial role in the regulation of histone deacetylation. This hypothesis is further supported by the result that key components of the NuRD complex, including Mbd3, Chd4, Mta1, Mta3, Rbbp4 and Rbbp7, maintained stable expression levels from the GV stage to the MI stage (Fig. 4f). Although the histone deacetylation is completed at the MI stage, the observation that Sin3a undergone translational activation at the MI stage and accumulated to high abundance at the MII stage (Fig. 4e, Supplementary Fig. S5), prompts further investigation into the potential role of the Sin3 complex in epigenetic regulation^50,51^.

### Phospholipid metabolism-mediated myristoylation modification closely associated with oocyte maturation

Cellular activities and metabolic pathways are regulated through complex, multilayered networks that integrate enzyme expression, metabolite dynamics, and PTM regulation. The scTPPM approach enables simultaneous profiling of mRNAs, proteins, metabolites, and PTMs at single-cell resolution, allowing integrated analysis of regulatory pathways from upstream metabolic enzymes to downstream PTM regulation. Using the phospholipid metabolic pathway as an example (Fig. 5a), the conversion of membrane phosphatidylcholine (PC) to protein myristoylation involves key enzymes, including Pla2g4c, Lypla1, Acsl6, and Nmt1, as well as intermediates such as lysophosphatidylcholine (LPC), choline glycerophosphate (GPC), long-chain fatty acids, and acyl-CoAs. Within the scTPPM framework, transcriptomic data capture mRNA expression of metabolic enzymes, proteomic data reflect their protein abundance, metabolomic data quantify pathway intermediates, and PTM-omic data resolve modification sites and modification levels. Together, these layers enable a comprehensive and integrated depiction of phospholipid metabolism and its role in regulating oocyte development.

**Fig. 5:**
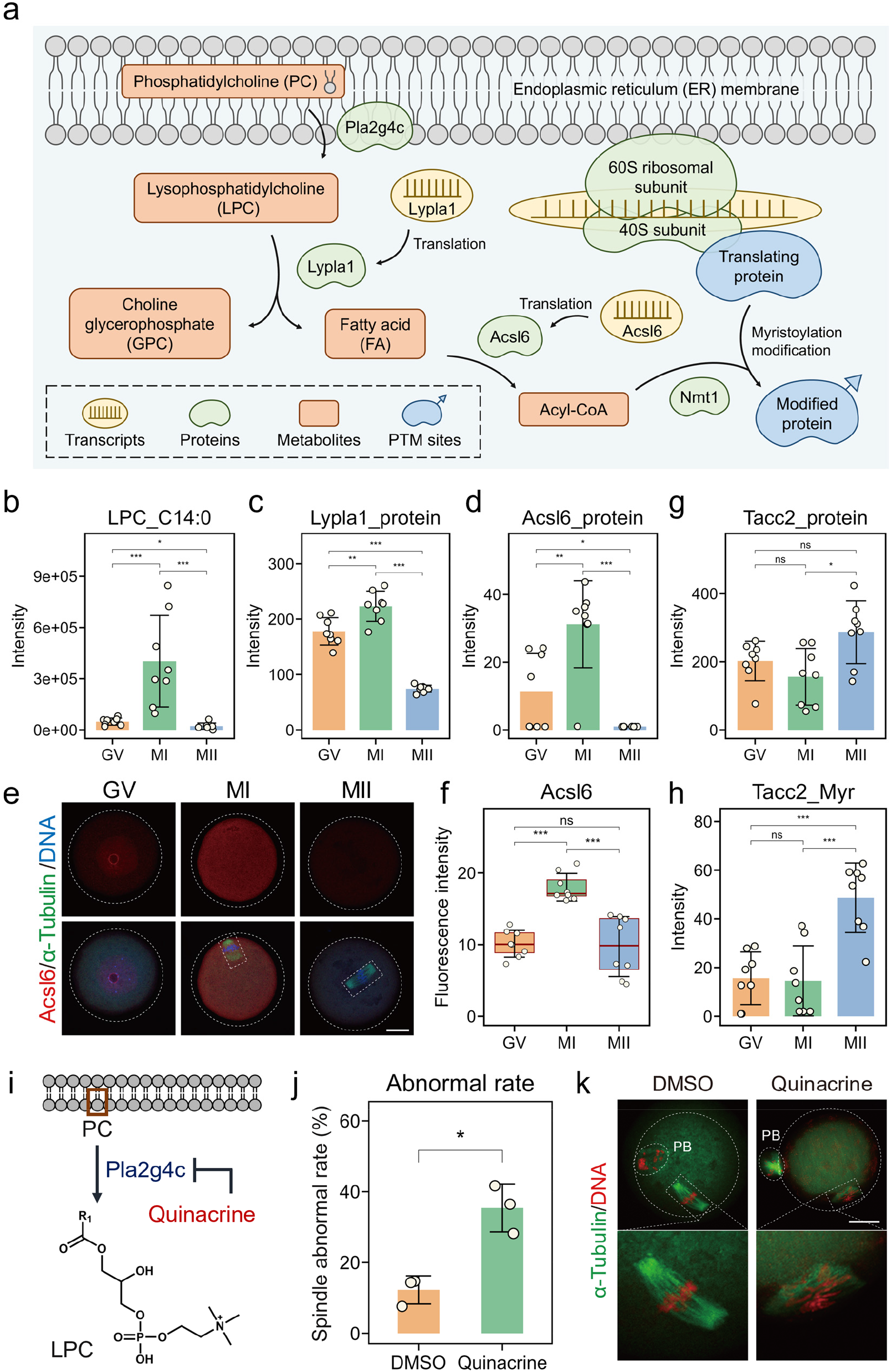
Phospholipid metabolism-mediated myristoylation modification of single mouse oocytes during meiosis maturation. a, Schematic diagram of phospholipid metabolism-mediated myristoylation from phospholipid choline (PC) on the membrane, through lysophosphatidylcholine (LPC), fatty acids (FA), and Acyl-CoA, to form myristoylation. b, Expression patterns of LPC (C14:0) in mouse oocytes at the GV (n = 8), MI (n = 8), and MII (n = 8) stages. c, Expression patterns of Lypla1 protein in mouse oocytes at the GV (n = 8), MI (n = 8), and MII (n = 8) stages. d, Expression patterns of Acsl6 protein in mouse oocytes at the GV (n = 8), MI (n = 8), and MII (n = 8) stages. e, Mouse oocytes at the GV, MI, and MII stages were immunostained on Acsl6 (Alexa Fluor 647, red), and α-tubulin (FITC, green), and fluorescently stained on DNA (DAPI, blue), scale bar: 25 μm. f, Fluorescent quantitative results of the oocytes immunostained on Acsl6 (Alexa Fluor 647, red) at the GV (n = 8), MI (n = 8), and MII (n = 8) stages. g, Expression patterns of Tacc2 protein in mouse oocytes at the GV (n = 8), MI (n = 8), and MII (n = 8) stages. h, Expression patterns of myristoylation of Tacc2 protein in mouse oocytes at the GV (n = 8), MI (n = 8), and MII (n = 8) stages. i, Schematic diagram of Quinacrine acting on the Pla2g4c protein to inhibit the hydrolysis of phospholipid choline (PC) to lysophosphatidylcholine (LPC). j, Abnormal rate of spindle in mouse oocytes treated with DMSO (n = 54) and Quinacrine (n = 62). k, Mouse MII oocytes treated with DMSO and Quinacrine were immunostained on α-tubulin (FITC, green), and DNA (PI, red), scale bar: 25 μm. *, p < 0.05; **, p < 0.01; ***, p < 0.001; ns, not significant; assessed by the student *t*-test.

Based on the multi-omics dataset, LPC (C14:0) was identified as a differentially expressed metabolite with a significantly increased abundance from the GV (n = 8) stage to the MI (n = 8) stage (Fig. 5b). This expression pattern was accompanied by the upregulated expression levels of its downstream degradation enzyme Lypla1 and Acsl6 during the GV-to-MI transition, indicating an upregulated LPC degradation (Fig. 5c-f). LPC (C14:0) is hydrolyzed by Lypla1 to release myristic acid, which is subsequently converted into myristoyl-CoA by Acsl6 and utilized by Nmt1for N-terminal proteins myristoylation. To explore the effect of increased LPC degradation on protein myristoylation, we established a targeted PTM library specifically to re-analyze the proteomic data for the myristoylation modifications using the customized PTM analysis workflow. Notably, we identified a myristoylation modification at the glycine residue on the N-terminal of Tacc2 protein, and the expression of this protein and its PTM was significantly upregulated from the MI stage to the MII stage (Fig. 5g, h). Given the role of Tacc2 in spindle assembly and chromosome segregation^52^, these results suggest that the LPC-derived myristoylation of Tacc2 contributes to oocyte maturation. To test this hypothesis, we treated GV-stage oocytes with a Pla2 inhibitor quinacrine to disrupt phospholipid metabolism (Fig. 5i). Quinacrine-treated oocytes exhibited a significantly higher incidence of spindle assembly defects during the oocyte meiosis (Fig. 5j, k), supporting a functional link between the phospholipid metabolism-driven myristoylation and spindle integrity. Together, this integrated analysis illustrates the utility of the scTPPM approach for resolving metabolic-PTM regulatory pathways and highlights the power of single-cell multi-omics in dissecting complex biological processes.

## Discussion

In the field of single-cell multi-omics analysis, the ongoing pursuit for technological advancement lies in achieving more dimensions, greater depth, and higher data quality. This study developed the scTPPM approach which enables simultaneous profiling of the transcriptome, proteome, PTMome, and metabolome within single-cell individuals, achieving unprecedented depth and breadth in single-cell multi-omics analysis. Leveraging the advantages of microfluidic techniques and nanoliter-scale reaction systems, combined with the technologies of next-generation sequencing and high-sensitive mass spectrometry, this approach enables the identification and quantification of tens of thousands of mRNAs, thousands of proteins and post-translational modification sites, and hundreds of metabolites from sub-cellular quantities of samples. In contrast to existing methods typically limited to dual-omics analyses (e.g., transcriptome-proteome or proteome-metabolome), our approach extends single-cell multi-omics analysis to not only transcriptomic, proteomic, and metabolomic data, but also PTM information, especially including histone modification data for epigenetic regulation analysis.

Compared to analysis of transcriptome, proteome, and metabolome, the simultaneous analysis of multi-PTMs within the same sample presents a greater technical challenge, which has not been achieved even in bulk samples with inputs below 10 µg^53,54^, while the protein quantity in single cells ranges from 0.05 to 50 ng. Currently, non-enrichment strategy and data-independent acquisition mode are typically employed for single-cell PTM analysis, and the raw data analysis including library-free or library-based workflow are primarily limited to three PTM types^33,34^. In contrast to the existing methods, the present laboratory-built library-based multi-PTMs profiling workflow enables mining different types of PTMs of interest from single-cell MS data without complex pretreatment or specific MS acquisition parameters. This workflow provides an elegant solution for achieving in-depth extraction of multiple PTMs information in data obtained by regular single-cell proteomic analysis, which were contained within the MS raw data but has remained largely unexplored. We propose that this workflow serves as an efficient tool for comprehensive profiling of PTMomes at the single-cell level, with broad application prospects in single-cell proteomics and multi-omics research.

To demonstrate the performance of the scTPPM approach within a specific biological process, we applied it in single-cell multi-omics profiling of mouse oocyte maturation. Based on the obtained single-cell multi-omics dataset encompassing the transcriptomes, proteomes, PTMomes, and metabolomes, we were able to profile oocyte maturation from multiple novel perspectives, including PTM regulation, epigenetic regulation, metabolic regulation, and temporal dynamic heterogeneity. Across different omics dimensions, the metabolic and PTM regulation exhibited higher activity than transcription and protein translation, whether among distinct maturation stages or across continuous timeline. This demonstrates the importance of analyzing metabolic and PTM regulation in revealing the complex dynamic characteristics of biological processes. Focusing on the phospholipid metabolic pathway, we conducted a comprehensive analysis of the entire chain from mRNAs and proteins to metabolites and PTMs, covering translational expression of metabolic enzymes, metabolic enzyme regulation, metabolite reactions, and PTM regulation across this entire process. This demonstrates the analytical depth and breadth of the scTPPM approach in profiling biological processes, as well as its substantial capability and potential to elevate multi-omics analysis from simple cell-type identification through cell clustering and differential compounds to a new perspective of comprehensive pathway analysis.

This study further evaluated the performance of the scTPPM approach for multi-omics characterization in single somatic cells, achieving a total identification and quantification of 9,592 genes, 2,519 proteins, 180 PTM sites, and 279 metabolites in single HeLa cells. This further demonstrates the broader application potential of the scTPPM approach, which is expected to address the urgent need for high-quality and in-depth integrated analysis of single-cell multi-omics data in studies of rare cells (e.g. CTCs), virtual cell models, and cellular interactions. By further enhancing the throughput of the scTPPM approach, we can expect to overcome limitations in its broader adoption within research that relies on large-scale data analysis. In summary, the development of the scTPPM approach enables simultaneous analysis of transcriptomics, proteomics, PTMomics, and metabolomics in single-cell individuals, marking a significant stride toward the ultimate aspiration of single-cell whole-omics analysis.

## Methods

### Cell culture

HeLa cell line was cultured in an incubator with 5% CO_2_ and 37 °C, using DMEM (Gibco) supplemented with 10% fetal bovine serum (FBS, Gibco) and 100 U/mL penicillin-streptomycin solution (Solarbio). Cells were detached using trypsin-EDTA solution (Jinuo) at 80% confluence, and washed twice with phosphate buffered saline (PBS, Gibco).

### Mouse oocyte collection

Mouse oocytes were collected from 6-8 week-old female C57BL/6 mice. All C57BL/6 mice were obtained from the Zhejiang Academy of Medical Sciences (the Laboratory Animal Center), and were housed under specific pathogen-free (SPF) conditions with 12 h light-dark cycles at 20–24 °C and 50%–70% humidity. All animal experiments were approved by the Zhejiang University Animal Research Ethics Committee (Ethics approval No. 21764). Female C57BL/6 mice were administered 10 IU of pregnant mare serum gonadotropin (PMSG, Sansheng) via intraperitoneal injection. After 44 h post-injection, the mice were euthanized and the ovaries were isolated. Fully grown GV stage oocytes were collected in M2 medium (Sigma-Aldrich). For in vitro maturation, the GV oocytes were cultured in M16 medium (Sigma-Aldrich) under mineral oil (Sigma-Aldrich) at 37°C in a 5% CO_2_ atmosphere. Oocytes typically reached the MI stage after 8 h and the MII stage after 16 h of culture.

### LC-MS/MS

In the proteomic analysis, the peptides were analyzed using an EASY-nLC 1200 LC system (Thermo Fisher Scientific) coupled with a timsTOF Pro mass spectrometer (Bruker Daltonics), using a home-packed nanoLC column (60 mm × 50 µm, 1.7 µm C_18_, 120 Å). The mobile phase A consisted of water with 0.1% formic acid and the mobile phase B consisted of 20% water in acetonitrile (ACN) with 0.1% formic acid. The flow rate of LC separations was 150 nL/min with a 70 min gradient as follows: 0–40 min, 0–40 % B; 40–41 min, 40-100 % B; and 41–70 min 100% B. The spray voltage of ion source was set at 1.75 kV under the positive mode. The acquisition mode of diaPASEF was used with a mass range of 350–1150 m/z, an isolation width of 25 m/z, mobility 1/k0 range of 0.6–1.3, and collision energy range of 20–47.3 eV in collision-induced dissociation (CID).

In the metabolomic analysis, the metabolites were also analyzed using an EASY-nLC 1200 LC system (Thermo Fisher Scientific) coupled with a timsTOF Pro mass spectrometer (Bruker Daltonics), using a home-packed nanoLC column (60 mm × 50 µm, 1.7 µm C_18_, 120 Å). The mobile phase A consisted of water with 0.1% formic acid and the mobile phase B consisted of 20% water in ACN with 0.1% formic acid. The flow rate of LC separations was 150 nL/min with a 70 min gradient as follows: 0–40 min, 0–40 % B; 40–41 min, 40-100 % B; and 41–70 min 100% B. The spray voltage of ion source was set at 1.75 kV under the positive mode. The acquisition mode of PASEF was used with a mass range of 100–1500 m/z, mobility 1/k0 range of 0.45–1.45, and collision energy range of 20–59 eV in CID.

### Raw data analysis

The raw data of single-cell mRNA sequencing were first trimmed using trim_galore (v 0.6.0) and validated using fastQC (v 0.12.1). The clean data were aligned with the reference genome (GRCh38 and GRCm39, https://ftp.ensembl.org) using hisat2 (v 2.2.1). The obtained sam files were converted into bam files using samtools (v 1.13). Based on the bam files, read counts of each sample were obtained through the featureCounts (v 2.0.1).

The raw data of single-cell proteomics and PTM profiles were analyzed using Spectronaut software (v 18.2.230802.50606, Biognosys) with library-based mode. The UniProt database (UP000000589.fasta, Mus musculus: 21,985 entries; UP000005640.fasta, Homo sapiens: 20,422 entries) was used in software analysis with the default settings. The false discovery rates (FDR) for both precursors and proteins were 0.01.

The raw data of single-cell metabolomics were analyzed using MetaboScape 2024b (v 13.0.3, Bruker Daltonics) with the T-ReX 4D workflow. In the peak detection parameters, the intensity threshold was set as 500 counts, and the minimum 4D peak size was set as 25 points. In the ion deconvolution parameters, the primary ion was set as [M+H]^+^ with the seed ions of [M+Na]^+^, [M+K]^+^, and [M+NH_4_]^+^. Spectral library was used for feature annotation, including Bruker MetaboBASE Personal Library (v 2.0 & v 3.0), HMDB (v 5.0), and MassBank (v 2024.11). For feature scoring and filtering, only features with MS/MS spectra were chosen and then filtered with the m/z tolerance of 15 ppm and the collision cross section (CCS) tolerance of 10%.

### Immunofluorescence assay

Oocytes were fixed with 4% paraformaldehyde-PBS solution (Sigma-Aldrich) for 30 min, permeabilized with 0.3% Triton X-100-PBS solution (Sigma-Aldrich) for 30 min, and then blocked in PBS containing 1% bovine serum albumin (BSA, Sangon Biotech) for 30 min. The oocytes were then incubated with primary and secondary antibodies (Supplemental Table S1) at 25°C for 1 h each. Nuclei were stained with 4′,6-diamidino-2-phenylindole (DAPI, Beyotime Biotechnology) for 15 to 30 min, and α-tubulin were then labeled with FITC conjugated anti-α-tubulin antibody (Sigma-Aldrich) for 30 to 60 min. Images were acquired using a Zeiss LSM900 confocal microscope (Zeiss).

### Inhibitor treatment

For inhibition experiments, oocytes were collected and pretreated with the Pla2 inhibitor quinacrine (Solarbio) for 6 h. Quinacrine was dissolved in DMSO (Sigma-Aldrich) with 4 mM stock solution and 10 μM working solution. Oocytes treated with DMSO were used as the control group. Following the quinacrine and the DMSO treatment, oocytes were continuously cultured in M16 medium (Sigma-Aldrich). The proportion of spindle abnormalities during meiosis maturation was quantified under different treatment conditions.

### Statistics and bioinformation analysis

Statistical significance was determined using the student *t*-test. The expression level of mRNA was represented using reads per million mapped reads (RPM) of each gene. The expression level of proteins was quantified using the default LFQ method in the Spectronaut software. The quantification value of peptides with target PTM sites was calculated using the total quantity of targeted EG in the Spectronaut software (v 18.2.230802.50606, Biognosys). The expression level of metabolites was represented using the integrated peak area in Metaboscape 2024b software (v 13.0.3, Bruker Daltonics). All statistical analyses and data visualizations were performed in R (v 4.4.2). Specifically, VennDiagram, UpsetR, and eulerr packages were used for analysis and visualization of set relationships. Seurat^55^ package was used to perform PCA, UMAP, and WNN analysis for clustering analysis and visualization. Mfuzz package was used for Fuzzy C-Means clustering analysis. ClusterProfiler package was used for GO analysis of target gene lists. Limma package was used for differential expression analysis of the transcriptome, proteome, PTMomes, and metabolome. Slingshot package was used to perform the pseudotime analysis. Pheatmap package was used for heatmap visualization. In addition, the three-dimensional structures of proteins and protein complexes were predicted using the AlphaFold3 online server (https://alphafoldserver.com).

## Acknowledgements

The authors acknowledge the financial support provided by the National Key R&D Program of China (Grants 2025YFA1309402, 2021YFA1301601, and 2024YFA1308400), the National Natural Science Foundation of China (Grants 22527805, 22404145, 22234007, and 82201837), the Zhejiang Provincial R&D Program (Grant 2024C03144), and the China Postdoctoral Science Foundation (Grant 2025M770968).

## Contributions

Q.F., X.D.F., and Y.R.J. conceptualized the study. Y.R.J., L.R.C., Y.M.Q., and Y.X.H. performed the experiments. Y.R.J., X.F.N., J.W., L.R.C., and Y.Y. performed the data analysis and generated the figures. Y.R.J., Q.Q.X., Q.W., Y.W., and J.Z.P contributed to optimizing and performing the equipment system. Y.R.J., L.R.C., and X.F.N. wrote the manuscript. Q.F., X.D.F., H.S.G., Y.R.J., L.R.C., and X.F.N. revised the manuscript. Q.F., X.D.F., H.S.G., J.Z.P., and Y.Y. contributed to the supervision of this work. All authors read, edited, and approved the final version of the manuscript.

