## Supporting information for "Single-cell Transcriptome-Proteome-PTMome(×8)-Metabolome (scTPPM) co-profiling decodes multimodal regulation during mouse oocyte meiosis"

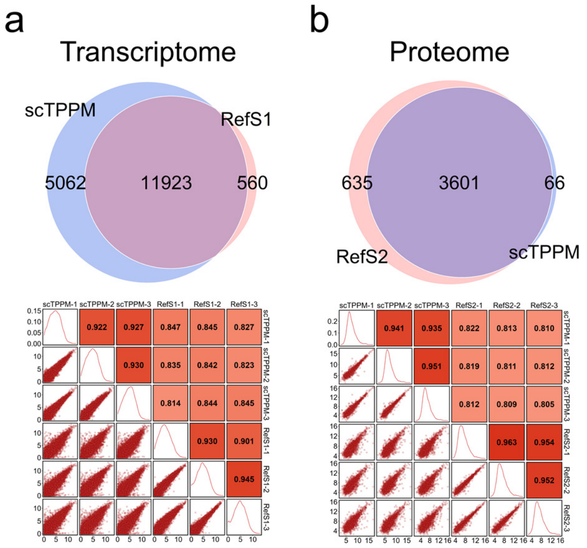


Supplementary Fig. S1: Comparative analysis between our results and the published results of single mouse oocytes. a, Venn diagrams and correlation analysis of the identified and quantified transcribed genes of the GV samples in our data and the published data of Ref. S1 (Zhang *et al.* 2025). b, Venn diagrams and correlation analysis of the identified and quantified proteins of the GV samples in our data and the published data of Ref. S2 (Zhang *et al*. 2025).

Reference S1: Zhang, J. *et al.* Vitrification preserves oocyte transcriptome in a 3D in vitro follicle development and oocyte maturation system. *Biol. Reprod.* **113**, 482-486 (2025).
Reference S2: Zhang, H. *et al*. Deep profiling of oocyte aging enabled by simple one-step vial-based pretreatment and single-cell proteomics. *JACS Au* **5**, 2321-2333 (2025).

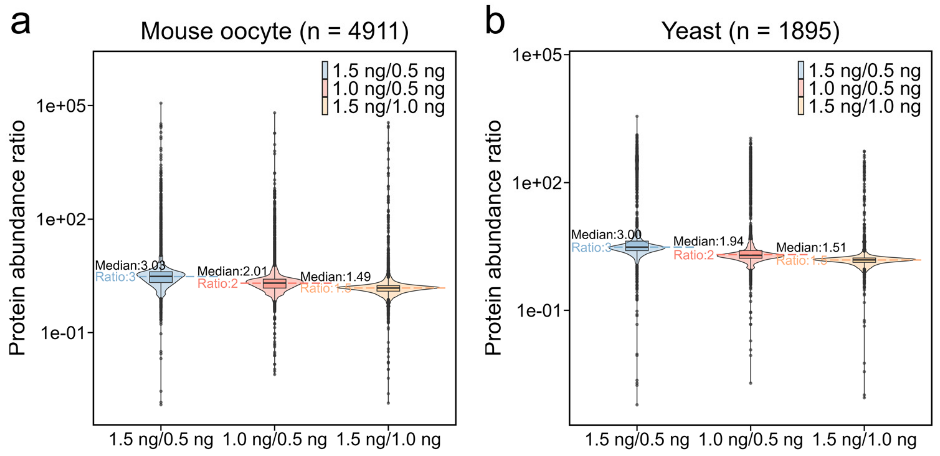


Supplementary Fig. S2: The benchmark analysis of oocyte (a) and yeast (b) quality control samples mixed in different proportions (25%, 50%, 75%).

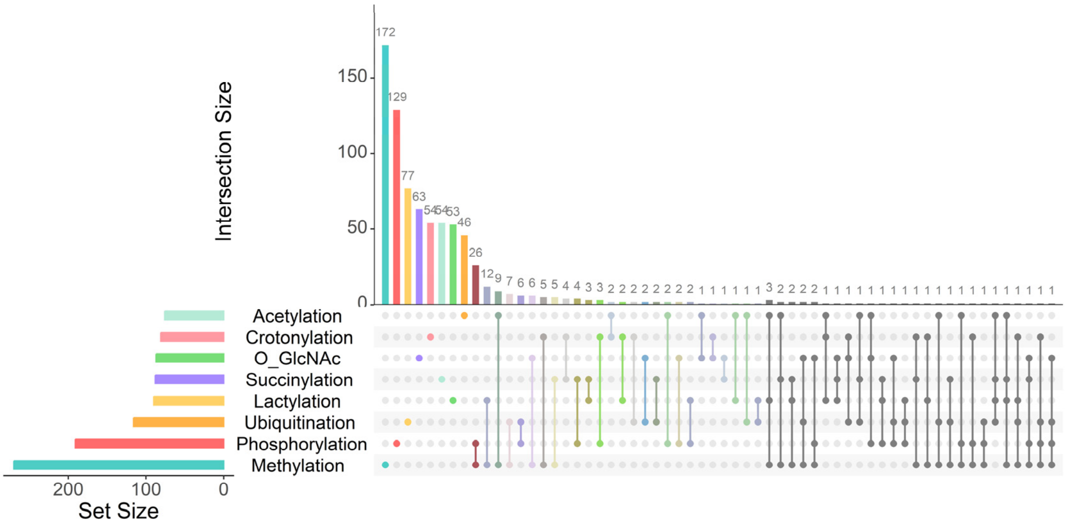


Supplementary Fig. S3: Set relationships among eight types of PTMs using Upset diagram.

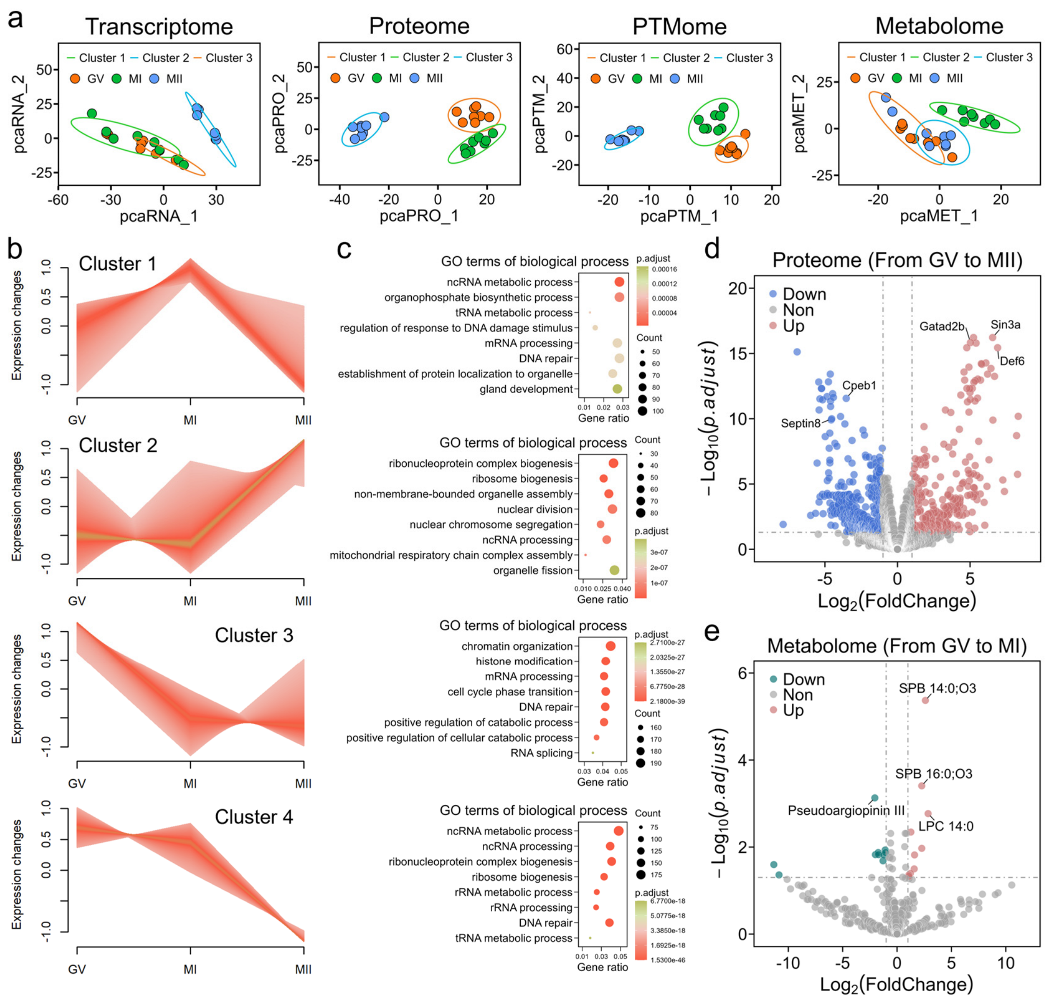


Supplementary Fig. S4: Clustering and differential expression analysis of single mouse oocytes during meiotic maturation. a, Visualization of PCA dimension reduction and KNN cluster analysis of single mouse oocytes at the GV, MI, and MII stages using transcriptome, proteome, PTMome, and metabolome dataset based on the scTPPM workflow. b, Fuzzy C-Means clustering of the transcriptomic data to categorize the genes into four clusters. c, GO enrichment analysis of the gene clusters identified by the Fuzzy C-Means clustering. d, Differential expression analysis of proteomic data in mouse oocytes from the GV stage to the MII stage. e, Differential expression analysis of metabolomic data in mouse oocytes from the GV stage to the MI stage.

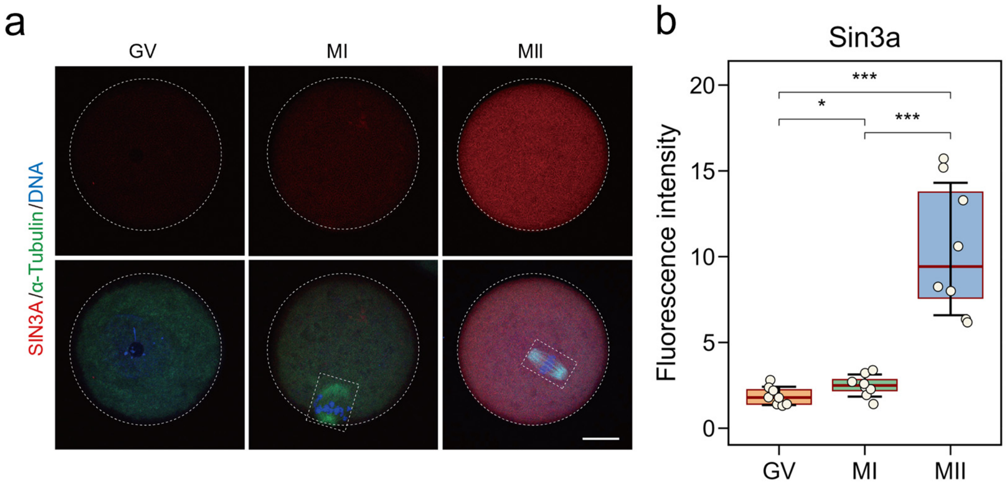


Supplementary Fig. S5: a, Mouse oocytes at the GV, MI, and MII stages were immunostained on Sin3a (Alexa Fluor 647, red) and α-tubulin (FITC, green), and fluorescently stained on DNA (DAPI, blue), scale bar: 25 μm. b, Fluorescent quantitative results of oocytes immunostained on Sin3a (Alexa Fluor 647, red) at the GV, MI, and MII stages.

Supplementary Table S1: List of antibodies for immunofluorescence analysis.

| Antibody | Conjugate | Company | Catalog No. |
|---|---|---|---|
| Rabbit mAb anti-Acetyl-Histone H4K12 | Unconjugated | Cell Signaling | #13944T |
| Rabbit mAb anti-Acetyl-Histone H4K16 | Unconjugated | Cell Signaling | #13534T |
| Rabbit mAb anti-Acetyl-Histone H3K23 | Unconjugated | Abcam | #ab177275 |
| Rabbit mAb anti-SIN3A | Unconjugated | Cell Signaling | #7691T |
| Rabbit pAb anti-LYPLA1 | Unconjugated | Thermo Fisher | #PA5-101405 |
| Rabbit pAb anti-ACSL6 | Unconjugated | Thermo Fisher | #PA5-101652 |
| Rabbit pAb anti-NMT1 | Unconjugated | Thermo Fisher | #PA5-78082 |
| Goat anti-rabbit IgG secondary antibody | Alexa Fluor 647 | Thermo Fisher | #A21244 |
